# Localization of genetic determinants for pathogenicity of Maize dwarf mosaic virus and Bermudagrass southern mosaic virus

**DOI:** 10.1101/651372

**Authors:** Farveh sadat Mostafavi Neyshabouri, Ahad Yamchi, Seyed Kazem Sabbagh, Mahmoud Masumi

## Abstract

Maize dwarf mosaic virus (MDMV) and Bermuda grass southern mosaic virus (BgSMV) are economically important potyviruses of cereals. BgSMV is very similar in genomic sequence to MDMV, but cannot infect Johnsongrass and is not transmitted by *Rhopalosiphum maidis*. Comparison of their genomes showed an additional stretch of 90 nucleotides in BgSMV coat protein but not in MDMV. Since the 90-nucleotide region is located in the N-terminal of BgSMV coat protein, it seems to have a role in biological properties such as vector transmission and pathogenicity. Recombinant virus constructs were made with and without the 90 nucleotides using SOEing PCR (MDMV (+90) and BgSMV (−90). Johnsongrass plants inoculated with the wild-type MDMV and recombinant BgSMV (−90) showed mosaic symptoms after 16 and 23 days, respectively, whereas plants inoculated with the wild-type BgSMV and recombinant MDMV (+90) didn’t show any symptoms until three months after inoculation. The qRT-PCR results detected significantly higher levels of BgSMV (−90) and MDMV compared to BgSMV and MDMV (+90), respectively. Also, *R. maidis* was able to transfer only the wild type MDMV and BgSMV (−90) from infected to healthy plants. These results confirmed that the insertion of the 90-nt region into the coat protein of MDMV affects the pathogenicity of the virus.

## Introduction

Potyviruses are the largest and economically the most important plant viruses. Virions are flexuous rods, 680–900 nm long × 11–15 nm wide. Their genome is a single-stranded RNA of positive polarity with a length of approximately 10 kb. The RNA genome carries a VPg (viral protein, genome-linked) covalently bound to its 5’ end, and a poly (A) tail at its 3’ end. The genome contains a single long open reading frame (ORF) that is proteolytically processed into proteins including P1, HC-Pro, P3, PIPO, 6K1, CI,6K2, VPg, NIa-Pro, Nlb, and coat protein (CP) by virus-encoded proteinases that are part of the polyprotein (Gibbs *et al*. 2010).

In Iran, Maize dwarf mosaic virus (MDMV) and Bermuda grass southern mosaic virus (BgSMV) are the most important cereal potyviruses (Mostafavi *et al*., 2015; Izadpanah *et al*., 2003). Weed plants of the cereal family, particularly the perennial species such as Bermuda grass are the host of a large number of plant viruses and some of them are economically important. Bermuda grass (*Cynodondactylon*) is widely disturbed in different climates in the world. Several viruses such as Barley yellow dwarf virus (BYDV), Bermuda grass etched-line virus (BELV), Cynodon chlorotic streak virus (CCSV), and Tomato spotted wilt virus (TSWV) has been reported from Bermuda grass in the world (Jorda*et al*., 1995; Lockhart *et al*., 1985 a, b; Louie 1980; Hosseini et al. 2010). In Iran, Bermuda grass is a source of several viruses (Izadpanah *et al.*, 2003). Studies showed that the viruses causing mosaic in Bermuda grass are classified into two groups; The first group includes isolates of *Bermuda grass southern mosaic virus* (BgSMV) which have a widespread distribution in tropical areas of southern Iran infect maize, Bermuda grass and goose grass with mosaic symptoms that its complete genome was sequenced (Zare et al. 2005). Based on molecular studies, BgSMV has the highest correlation with MDMV, sorghum mosaic virus (SrMV) and sugarcane mosaic virus (SCMV). The second group consists of isolates of Bermuda grass mosaic virus (BgMV).

MDMV is one of the most common viruses with a worldwide distribution and causes severe losses in maize and sorghum crops (Uzarowska *et al*. 2009). MDMV and BgSMV are transmitted via mechanical inoculation and aphids in a non-persistent manner (Leng *et al*. 2015). The results of the study on serology, sequencing, host range and vector transmission showed that BgSMV is similar to MDMV but is different in three properties: all of BgSMV isolates available in GenBank have an additional 90-nucleotide region compared to MDMV in 5’-region of the CP gene. In contrast to MDMV, BgSMV is not transmitted by *Rhopalosiphum maidis* aphid and cannot infect Johnsongrass which is the primary inoculum source or original host of MDMV (Zare *et al.*, 2005).

The 90-nt region is located in the same nucleotide position (nucleotide 108-198) of the CP of the isolates of BgSMV. This region is highly conserved (nearly 100%) in all known BgSMV isolates but there is no information about the function of this 90-nt region.

To understand the potential role of this region in non-pathogenic BgSMV, and its potential role in transmission by *R. maidis*, we made infectious transcripts of recombinant constructs with and without the 90-nt region to study the role of this region in viral pathgenecity and vector transmission.

## Results

The MD3F/MD1R primer, designed for the flanking region of the 90-nt, amplified 636 bp and 726 bp fragments from MDMV and BgSMV genomes, respectively (Fig3). SOEing PCR results showed the correct join of MD1, MD2 and MD3 fragments by overlapping region among these fragments (fig 5B-C) and fig 5A depicts the assembling of Bg1 and Bg2 fragments based on related its overlapping. Finally, the recombinant MDMV (+90) and BgSMV (−90) were produced. The fig 5D and E shows that we could add successfully T7 promoter and poly A tail to wild type MDMV and BgSMV.

The Johnsongrass plants inoculated with the wild-type MDMV and recombinant BgSMV (−90) showed mosaic symptoms after 16 and 23 days, respectively, whereas the same plants inoculated by the wild-type BgSMV and recombinant MDMV (+90) didn’t show any symptoms until three months after inoculation (Fig 6A-C). Also, Sequencing result of the RT-PCR product derived from infected Johnsongrass (BgSMV (−90)) showed the multiplication of the virus which lacks the expected 90-nt (Fig. 6D).

### Titer of viruses in inoculated plants

The standard curve, obtained by serial dilution series of recombinant plasmid carrying consensus segment of P1 gene of virus (refer to Fig 2), showed a linear relationship with high correlation (R^2^ = 0.994) between the amount of input DNA and the threshold cycle values (Ct) for over a range of seven log units (Fig 7). This curve was used as a standard reference for the extrapolation of copy number of virus targets of unknown concentrations in Johnsongrass plants. The number of virus copies/mg of tissue was in the range of 4.6×10^2^,4.7×10^5^,4.9×10^7^and 8.3×10^2^ in BgSMV, BgSMV (−90), MDMV and MDMV (+90) respectively with significant differences among them (p-value ≤ 0.01).

### Aphid transmission

*Rhopalosiphum maidis* was able to transfer only wild-type MDMV and BgSMV (−90) from inoculated to healthy plants. The mosaic symptoms of the disease were observed in plants fed with the infected aphid. Meanwhile, wild-type BgSMV and MDMV (+90) were not transmitted by *R. maidis* to healthy plants and no symptoms were observed. The results verified by RT-PCR using specific primer MDF3/MDR1 (Fig 8).

## Discussion

Bermuda grass southern mosaic virus (BgSMV) is very much similar in genomic sequence to Maize dwarf mosaic virus (MDMV), but does not infect Johnsongrass. Comparison of their genomes showed an extra fragment of 90-nt in BgSMV coat protein but not in MDMV. In this research, the potential role of the 90-nt region in the pathogenicity of the viruses in Johnsongrass was studied. Since the presence of additional 90-nt located in N-terminal of coat protein, it seems that it affects biological properties such as transmission with *R. maidis* and pathogenicity of the viruses. So, we made the recombinant viruses with and without the 90-nt using SOEing PCR method without cloning in expression vectors because of the large length of the virus genome which made limitation of cloning in expression vectors. Then, the sequence of T7 promoter and poly (A) tail was designed in recombinant and wild-type virus genomes through SOEing PCR in order to facilitate the *in vitro* transcription of the viruses.

The poly (A) tail at the 3’ end of the genome of some viruses have a protective role against exoribonuclease and it is involved in translational regulation and the synthesis of the negative viral RNA (Bleeman and Parker, 1995). Studies involving viral RNAs transcribed *in vitro* showed the effect of poly (A) tail length on virus infectivity (Guilford 1991). Tatineni *et al*. (2015) designed the infectious cDNA clone of *Triticum* mosaic virus to transcribe virus with 105 A tail residues. In another study, the use of a short poly (A) tail of 21 nucleotides reduced the pathogenicity of MDMV (Stewart *et al*. 2012). The white clover mosaic virus transcript without a poly(A) tail at the 3’ end decreased 50-fold symptom lesions lower than a transcript with 74 A residues, whereas a transcript with 10 A residues produced only twofold fewer lesions (Guilford 1991). Tacahashi and Uyeda 1999 made clover yellow vein virus synthetically with different poly A tail from 5 to 60 nucleotides and reported that the minimum length of poly A for pathogenicity is 5-10 nucleotides. Therefore, in this study, the length of the T tail with 58 nucleotides was designed in the reverse primer.

The RNA genome of potyviruses carries a VPg (viral protein genome linked) covalently bound to its 5’ end. This protein acts as a primer for complementary viral RNA synthesis (Jiang and Laliberte 2011). In the study of Stewart et al. (2012), despite the use of the cap analog in *in vitro* transcription reaction of MDMV, there was no significant pathogenicity after the injection of RNA synthesized into maize seeds. It is likely that the presence of the artificial cap simultaneously with VPg in virus genome by *in vitro* transcription may affect negatively the pathogenicity. Therefore, we did not use the cap or its analogues in the *in vitro* transcription reaction, because the wild-type viruses are cap-free. Pathogenicity was occurred by observing symptoms and confirming the presence of many copies of the pathogenic virus in susceptible plants (fig 6A-C).

The detection of MDMV (+90) and BgSMV (−90) from the infected plants were verified by RT-PCR until 23dpi. But only the plants infected by BgSMV (−90) and wild type MDMV showed the positive result in the RT-PCR. It can be interpreted that the RT-PCR detects the *in vitro* transcribed raw recombinant MDMV (+90) RNA from inoculated plants until 23dpi. Also, MDMV (+90) unlike BgSMV (−90) could not able to infect the plant because of the extra 90-nt. Furthermore, the absolute real time PCR calculated the concentration of MDMV (+90) RNA at Ct value 36 versus BgSMV (−90) RNA at Ct value 25 that verified the aforementioned RT-PCR result. In addition, it can be concluded that the RNA of both recombinant constructs was stable after inoculation, so they could be detected up to 23dpi. In order to analyze accurately the pathogenicity, we used the same concentration of recombinant viral RNA for inoculation of the plants.

In BgSMV and MDMV, phylogenetic analysis showed that BgSMV is the closest virus to MDMV and probably MDMV and BgSMV have diverged from each other (Zare et al. 2005; Hosseini et al. 2010; Mostafavi et al. 2015). Also, the ratio (ω) of non-synonymous (amino acid change) to synonymous (silent) substitutions which provide an estimate of the selective pressure at the protein level obtained low value for both of BgSMV and MDMV (Wolf et al. 2009; Hosseini et al. 2010; Mostafavi et al. 2015). Thus, purifying selection for MDMV and BgSMV population during evolution suggest that the deletion of the 90-nt from MDMV may has occurred to adapt with different conditions along with its pathogenicity on the Johnsongrass. Likewise, distribution of BgSMV in the southern regions of Iran, despite the many natural barriers, indicates the long history of this native virus in Iran.

The DAG conserved motif in the coat protein of potyviruses affects the transmission of the virus by aphids, locates upstream of the 90-nt region in BgSMV (Aterya *et al*. 1991; Lopez-Moya et al. 1999). Therefore, we guess that it is likely, the 90-nt affect the context of the motif and subsequently virus transmission negatively, but to prove this, it is necessary to check the interaction of aphid-virus proteins.

## Material and methods

### Source of viruses

Johnsongrass plants separately infected with Maize dwarf mosaic virus and Bermuda grass southern mosaic virus were maintained under controlled greenhouse conditions at the Plant Virology Research Center, Shiraz University, Iran. Since the viruses could not be distinguished by ELISA, virus-specific primers were used in RT-PCR (MD3F / MD1R) flanking the 90-nt region.

### Virus recombinant constructs

We used SOEing PCR to add the 90-nt from BgSMV to its corresponding region in MDMV which was referred to as MDMV (+90). Three overlapping primers with the 90-nt were designed with Oligo7 software to produce three fragments by RT-PCR (Fig 1 and Table 1). Fragments MD1 (8530bp) and MD3 (1079bp) were amplified by Expand Long Template PCR System (Roche, Germany) from MDMV genome using M1F/M1R and M3F/M3R, respectively. Fragment MD2 (107bp) with the 90-nt region was amplified from BgSMV genome using M2F/ M2R.

**Table 1.**
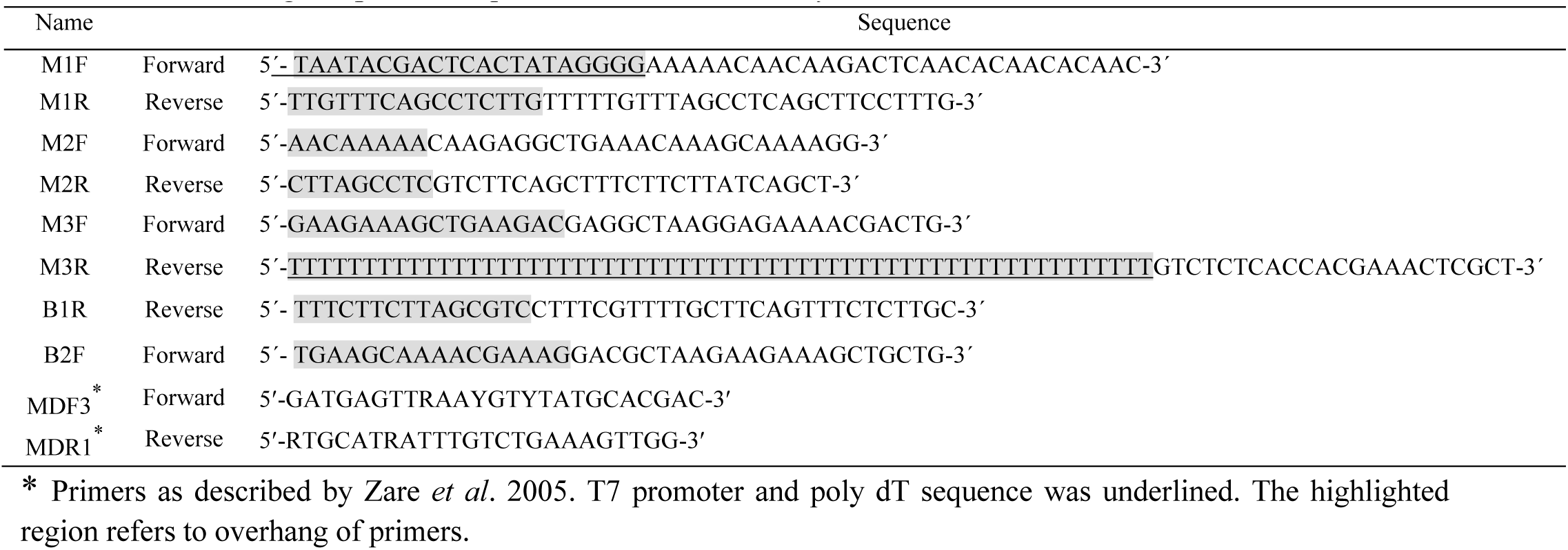
List of designed primer sequences used in this study

**Fig 1.**
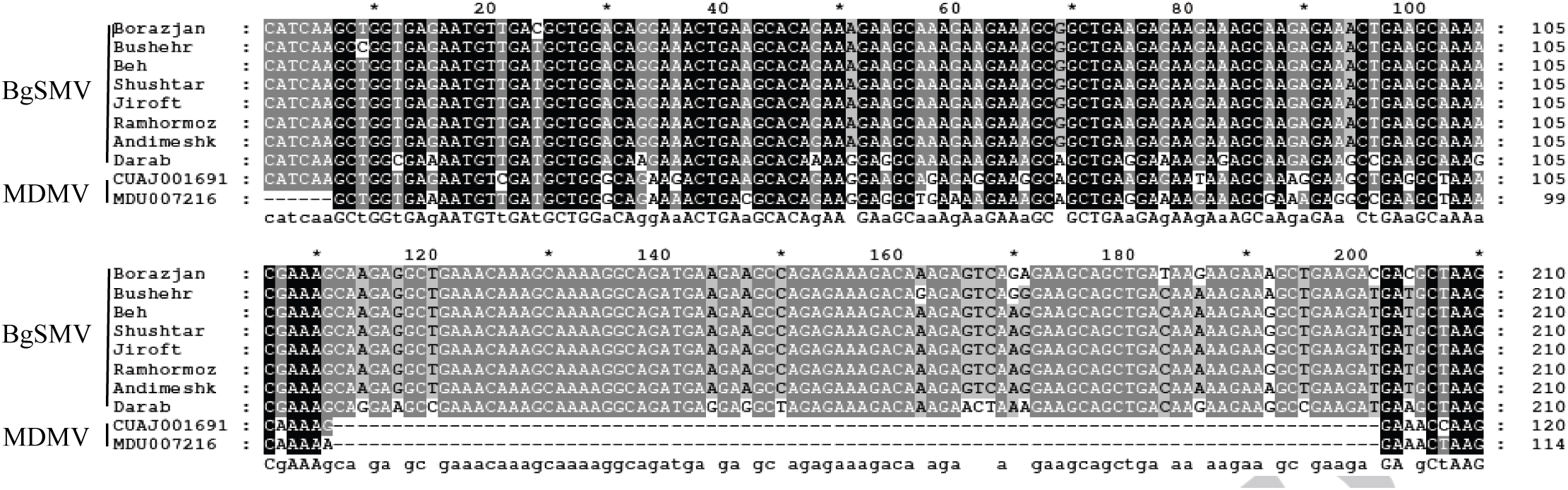
5’-region alignment for coat protein gene of Bermudagrass southern mosaic virus (BgSMV) and Maize dwarf mosaic virus (MDMV) isolates. The deleted 90-nucleotide region was shown at positions 112 to 201 corresponding to BgSMV isolates (Farahbakhsh *et al*., 2013).

**Fig 2.**
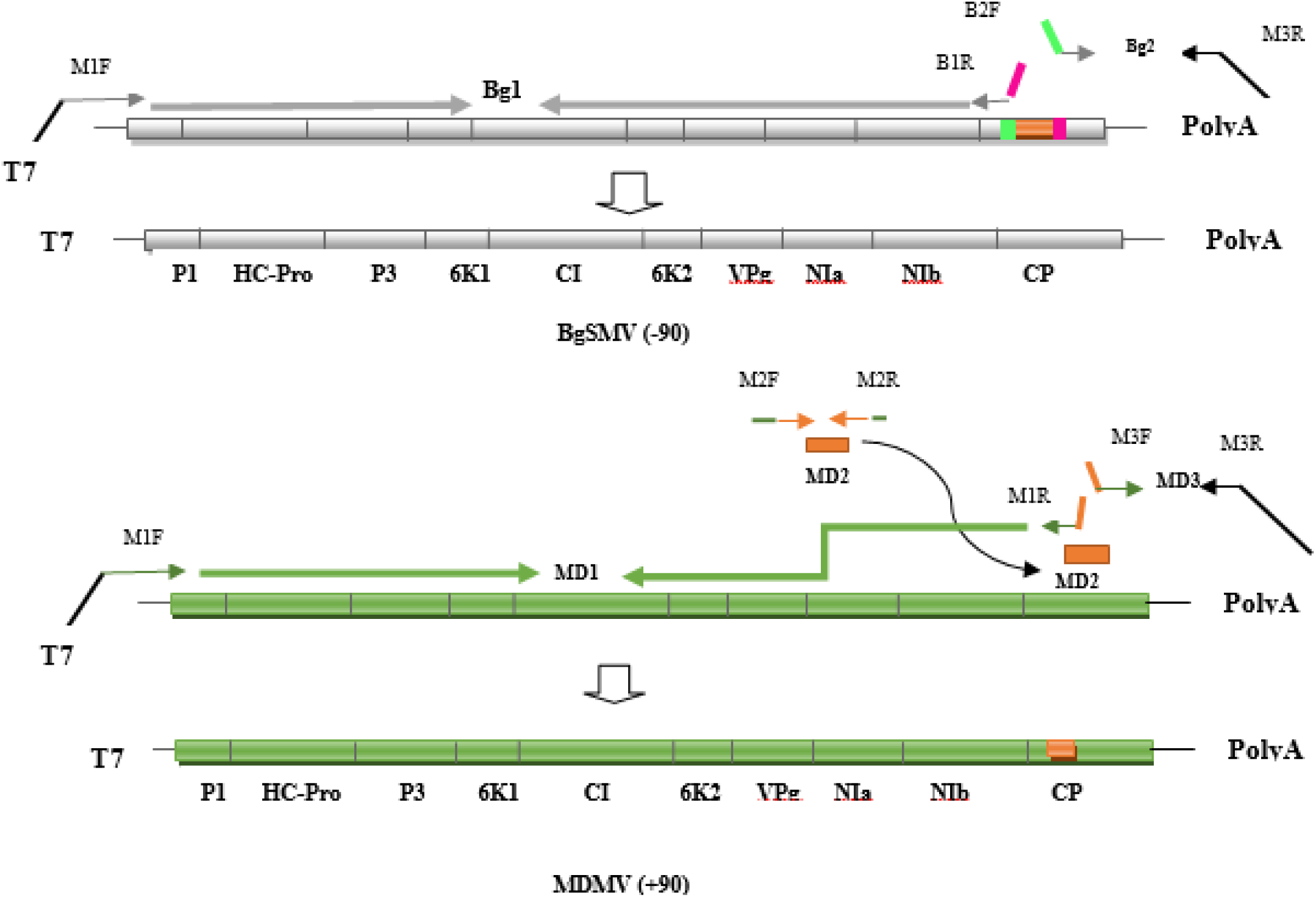
Position of primers on MDMV and BgSMV genomes

**Fig 3.**
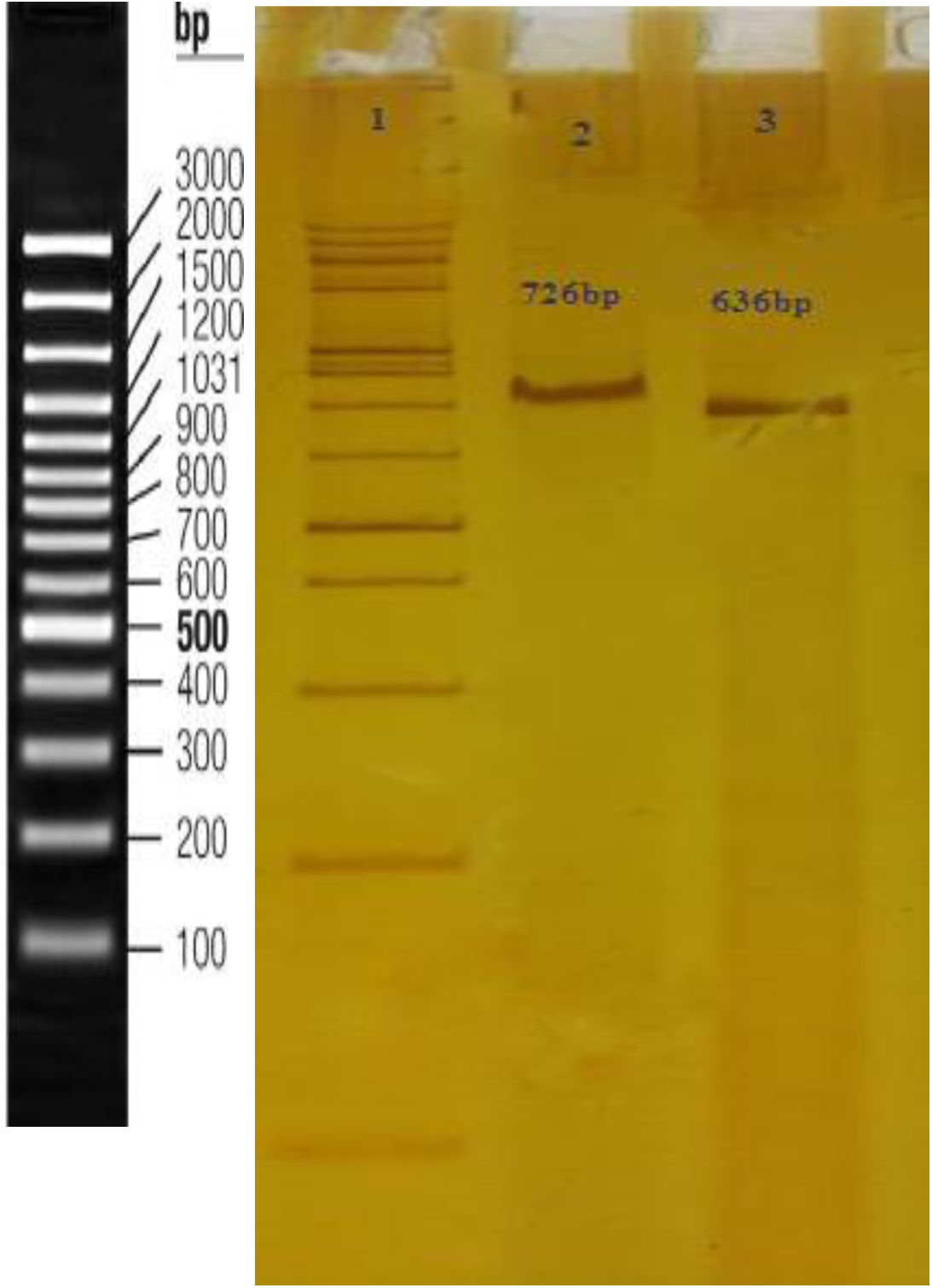
Electrophoresis on acrylamide gel; amplification of 636 bp and 726 bp fragments on MDMV and BgSMV by MD3F/MD1R primer, designed for flanking region of the 90-nt. Lane1. DNA ladder 100 bp+3K (SMOBIO); lane 2, BgSMV PCR product; lane3, MDMV PCR product.

**Fig 4.**
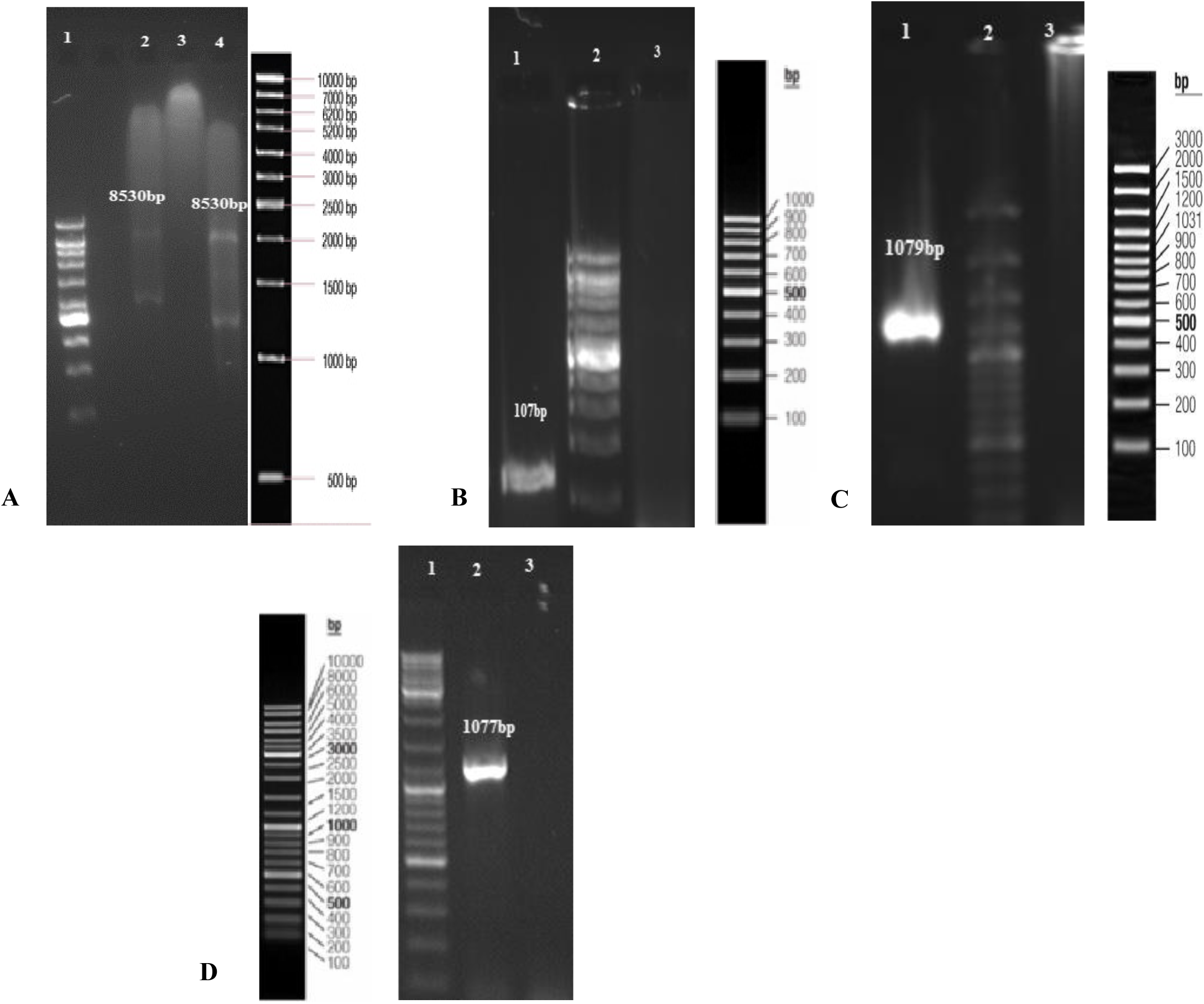
Electrophoretic analysis of the PCR products in 0.5% agarose gel. (A) Lane 1, DNA ladder 1KB (Vivantis); lane, 2 Bg1 fragment; lane 3, negative control; lane 4, MD1 fragment. (B) Lane 1, MD2 fragment; lane 2, DNA ladder 100bp (Fermentas); lane 3, negative control. (C) Lane 1, MD3 fragment; lane 2, DNA ladder 100 bp+3K (SMOBIO); lane 3, negative control. (D) Lane 1, Bg2 fragment; lane 2, DNA ladder 1KB (Vivantis); lane 3, negative control.

**Fig 5.**
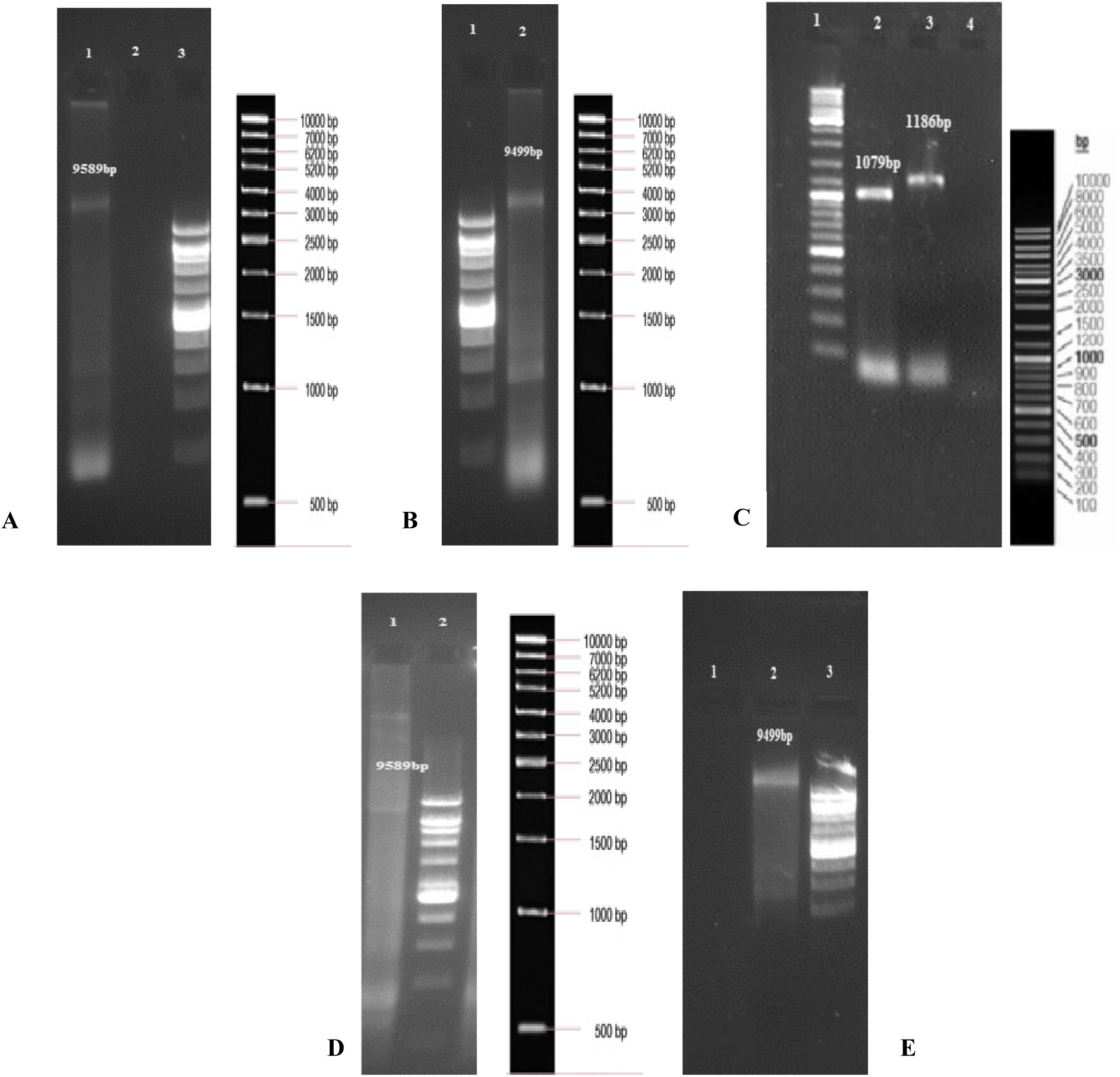
Electrophoretic analysis of the SOEingPCR products in 1% agarose gel. (A) Lane 1, MD1+(MD2+MD3) fragments; lane, 2 negative control; lane 3, DNA ladder 1kb (Vivantis). (B) Lane 1, DNA ladder 1kb (Vivantis); lane 2, Bg1+ Bg2 fragments. (C) Lane 1, DNA ladder mix (Fermentas); lane 2, MD3 fragment; lane 3, MD2+MD3 fragments; lane 4, negative control. (D) Lane 1, BgSMV-wild type, complete genome; lane 2, DNA ladder 1kb (Vivantis). (E) Lane 1, negative control; lane 2, MDMV-wild type, complete genome; lane 3, DNA ladder 1kb (Vivantis).

**Fig 6.**
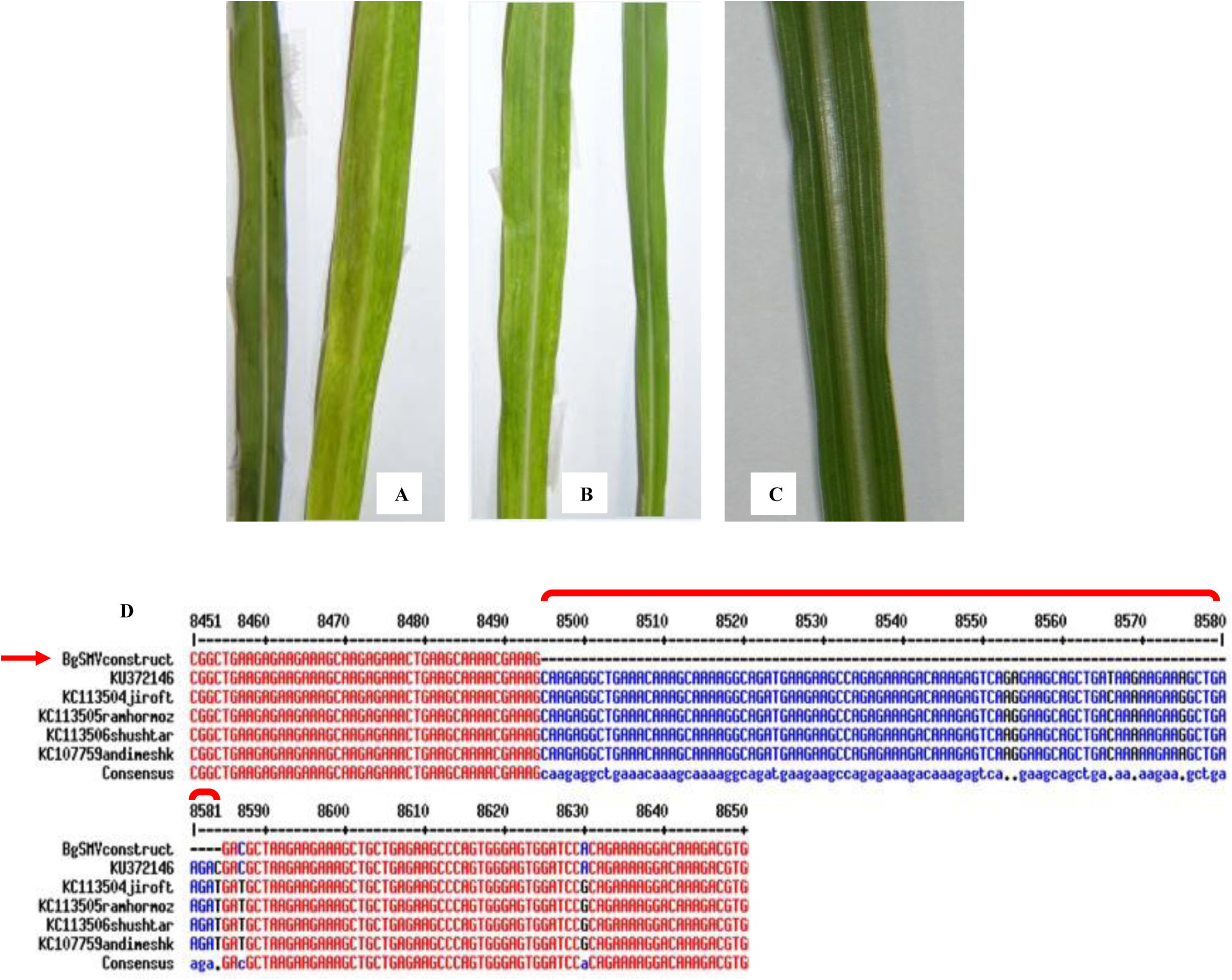
Johnsongrass plants inoculated by the wild-type and recombinant viruses; (A) No symptom of BgSMV wild type (left) and typical MDMV-wild type mosaic was observed in inoculated Johnsongrass (right). (B) Typical BgSMV (−90) mosaic (left) and no symptom of MDMV (+90) was observed in inoculated Johnsongrass (right). (C) Healthy leaf mock inoculated. (D) Sequencing result of RT-PCR product derived from infected Johnsongrass (BgSMV (−90)) was aligned with GenBank BgSMV wild type isolates. The bracket refers to the 90-nt deleted region from *in vitro* synthesized BgSMV.

**Fig 7.**
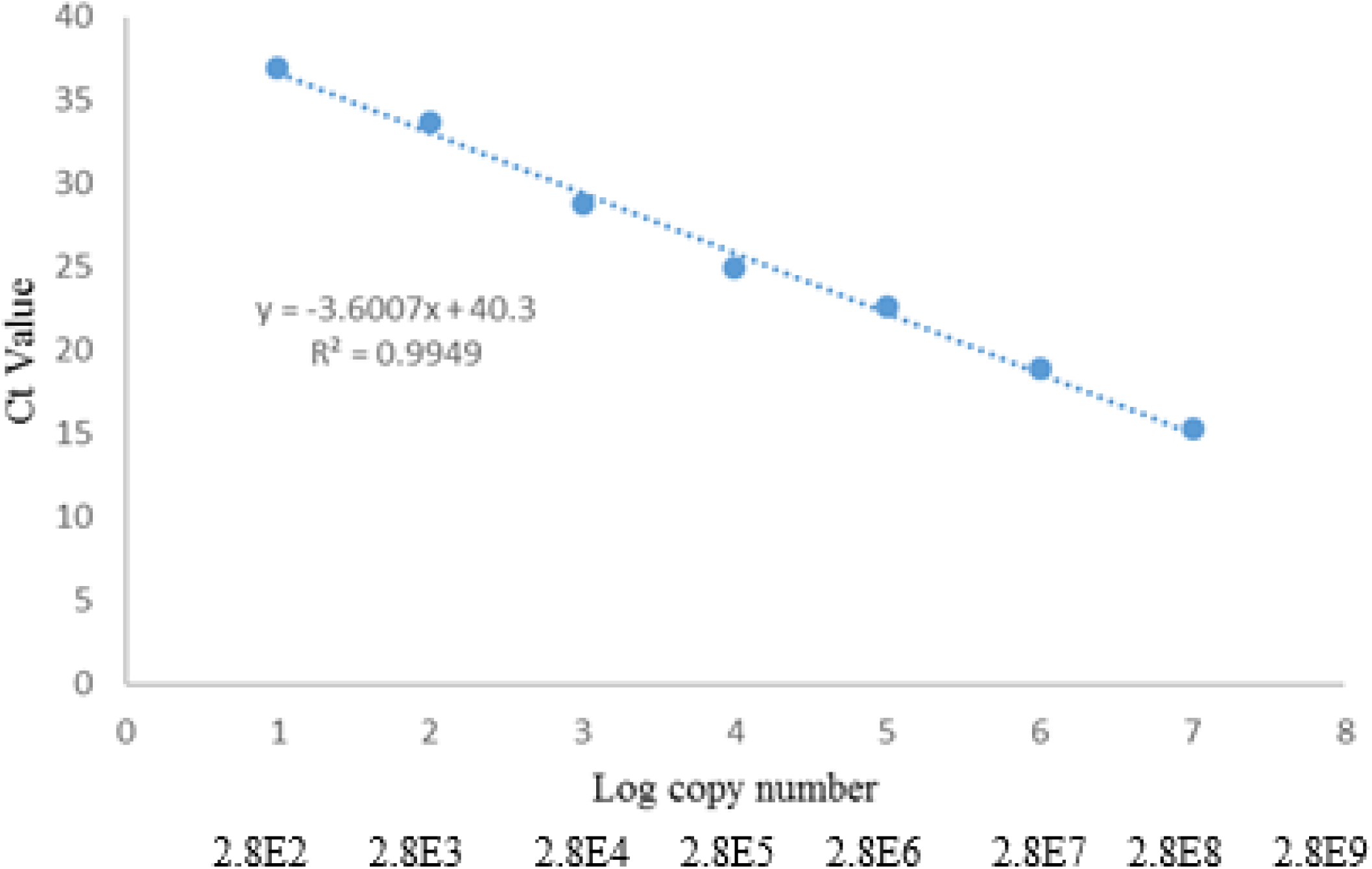
Linear range of the wild type viruses and recombinant constructs detection using SYBER Green absolute qPCR. Standard curve obtained by plotting of mean threshold cycle values over a range of 10-fold serial dilutions of the recombinant plasmid includes P1 segment. The horizontal axis reports the logarithm of number of copies of plasmid/μl while the vertical axis shows the threshold cycle values obtained in the reactions. The qRT-PCR was performed by four biological and three technical replicates.

**Fig 8.**
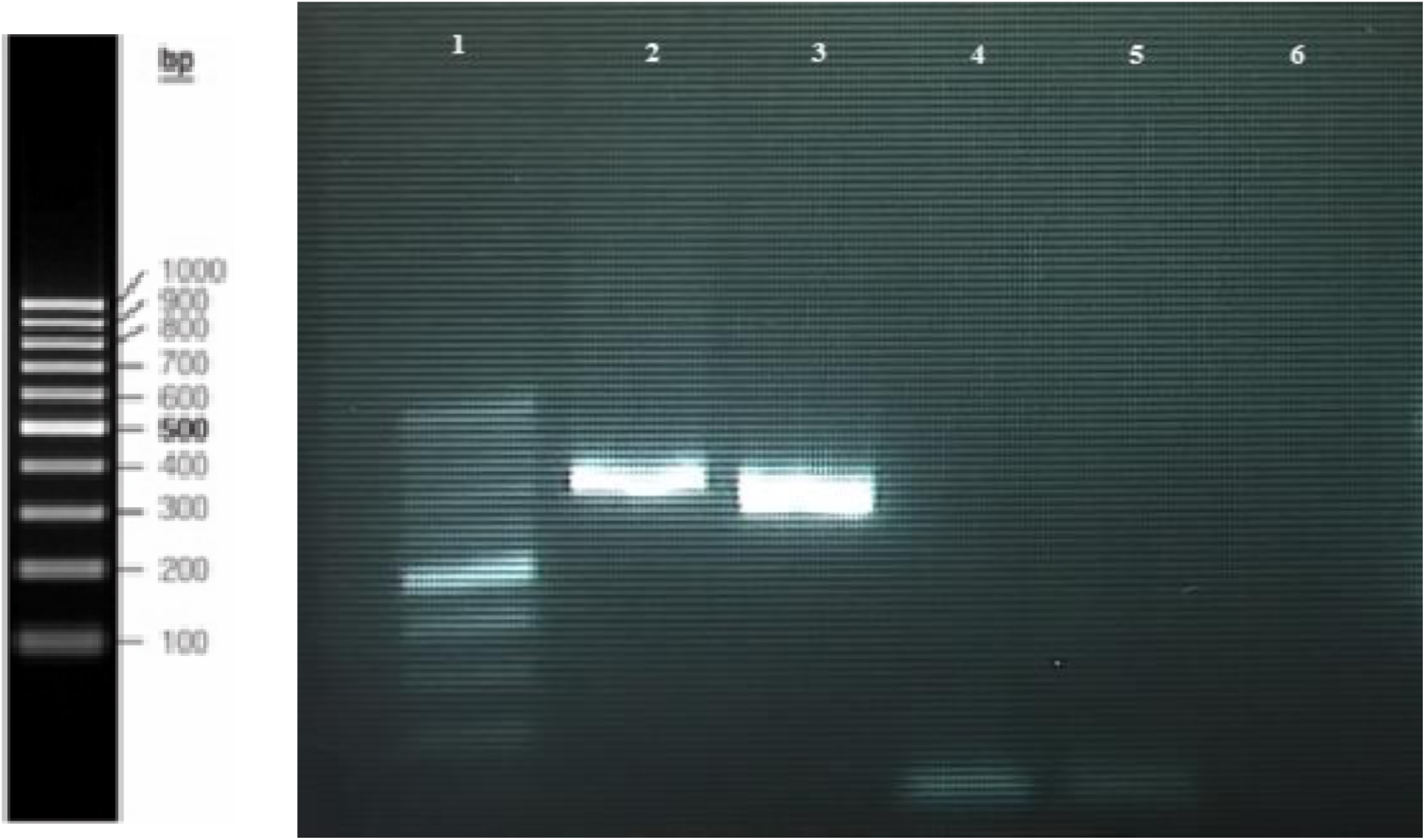
RT-PCR products of infected plants which fed by *R. maidis* carrying wild type and recombinant viruses; Lane 1, DNA ladder 100bp; lane 2, infected plant by *R. maidis* carrying wild-type MDMV; lane 3, infected plant by *R. maidis* carrying BgSMV (−90); lane 4, inoculated plant by *R. maidis* carrying wild-type BgSMV; lane 5, infected plant by *R. maidis* carrying MDMV (+90), lane 6, healthy plant.

To remove the 90-nt from BgSMV (BgSMV (−90), two overlapping primers lacking the 90-nt were designed with Oligo7 software. Two fragments, Bg1-8530bp and Bg2-1061bp, were amplified from BgSMV genome in PCR using M1F /B1R and B2F/ M3R (Fig 1 and Table 1). T7 promoter sequence and Poly T tails length of 58 nucleotides was embedded to the 5′-end and the 3′-end of the both constructs.

PCRs were individually purified using Qiaquick PCR purification kit (Qiagen). The purified products were then analyzed by electrophoresis in 1% precast agarose gels (Sigma).

Gibson assembly method (2011) was used to make MDMV (+90) and BgSMV (−90) constructs. Three MD1, MD2, and MD3 fragments were joined to create recombinant MDMV (+90) in two steps, A and B. In step A, the equimolar mixture of the MD2 and MD3 fragments was cycled 10 times without primers, and in step B a half of step A reaction was run with primers to obtain (MD2+MD3) fragment in 30 cycles. Then, the same as above A and B steps was done to join (MD2+MD3) fragment to MD1 to obtain the final recombinant MDMV (+90). To create of BgSMV (−90) construct, two Bg1 and Bg2 fragments were assembled in step A and B as described above.

step A: 5 µl PCR buffer, 2.5 µl 10 mM dNTP mix,5 µl MD2 fragment (60 ng), 12 µl MD3 fragment (100ng) to obtain MD2+MD3 and or 15 µl MD1 fragment (80 ng), 20 µl MD2+MD3 fragment (100ng) to obtain MD1+ (MD2+MD3) and 5 µl Bg1 fragment (60 ng), 10 µl Bg2 fragment (100ng) to obtain Bg1+Bg2, no primers, 0.75 µl Expand Long Template enzyme mix (Roche) bring to 50 µL with ddH^2^O, 50 µl of mineral oil on top after the ingredients are carefully mixed by pipetting with a wide-nozzled tip. Cycling parameters: to join MD2 and MD3 fragments, initial denaturation 94°C 2 min, subsequent steps 94°C 30 s, annealing 63°C 30 s, extension 68°C 1 min, 10 cycles total, final extension 1 min, hold at 4°C and to join MD1 and MD2+MD3 and or Bg1 and Bg2 fragments, 94°C 1 min, 0.23 °C/s to 25 °C, 25 °C 2 s, 50 °C 1 min, 0.13 °C/s to 68 °C, 68 °C 50 min, and then hold at 4°C. The resulting PCR was purified using Qiaquick PCR purification kit. About one-third of this DNA (10 µl) was used in the next step.

step B: 29.25 µl water, 5 µl PCR buffer, 2.5 µl 10 mM dNTP mix,10 µl of purified DNA from step A, 1.25 µl left primer (M1F or M2F or M3F or B2F), 1.25 µl right primer (M1R or M2R or M3R or B1R), 0.75 µl Expand Long Template enzyme mix (Roche), 50 µl of mineral oil on top after the ingredients were carefully mixed by pipetting with a wide-nozzled tip. Cycling parameters to join MD2 to MD3 fragments were: initial denaturation at 94°C for 2 min, subsequent steps at 94°C for 30 s, annealing at 65°C for 30 s, extension at 68°C for 1 min, 30 cycles total, final extension for 5 min, hold at 4°C. Cycling parameters to join MD1 to MD2+MD3 and Bg1 to Bg2 fragments were: initial denaturation at 94°C for 2 min, and for 15 cycles at 94°C for 10 s, annealing at 55°C for 1 min, extension at 68°C for 10 min, subsequent steps at 94°C for 15 s, annealing at 55°C for 1 min, extension at 68°C for 10 min plus 10 s per cycle, total 25 cycles, final extension for 10 min, hold at 4°C.

The M1F (containing T7 sequence) and M3R (containing poly T) primers were fused to wild type of MDMV and BgSMV genomes by PCR in order to carry out *in vitro* transcription. Likewise, these primers added T7 promoter and poly T tail to the recombinant MDMV (+90) and BgSMV (−90) to use for *in vitro* transcription reaction.

### *In vitro* transcription

For production of *in vitro* viral RNA transcripts, one µg of the recombinant constructs including MDMV (+90), BgSMV (−90), and their wild type viruses were used in the *in vitro* transcription reaction using T7 RNA polymerase. The resulting RNA transcripts were treated with DnaseI, RNase-free (Roche, Germany). A 1.5 µg of RNA transcripts were diluted in the 440 µl FES inoculation buffer (100ml GP buffer: 18.77gr glycine and 26.13 K2HPO4 bring to 500 mL with ddiH2O; 5gr sodium pyrophosphate, 5gr bentonite, 5gr celite bring to 500 mL with ddiH2O) and then mechanically inoculated to Johnsongrass at the two-leaf stage. Then, inoculated plants were maintained in a separate greenhouse under controlled conditions at 28 °C with 16h light/ 8h dark.

### Assessment of pathogenicity of *in vitro* transcripts on Johnsongrass

The inoculated Johnsongrass plants by recombinant constructs were evaluated based on mosaic symptoms, sequencing and absolute real time PCR.

Leaves were collected from Johnsongrass plants inoculated with RNA transcripts derived from recombinant constructs as well as wild type viruses. RT-PCR was carried out using specific primers (MDF3/MDR1). Sequencing was done on BgSMV-90nt and MDMV+90nt constructs.

For the purpose of absolute quantification, standard curves at a 10-fold serial dilution series of the PTZ57R plasmid carrying specific conserved fragment of the both viruses were prepared. The virus copies numbers of recombinant constructs as well as wild type viruses were calculated using the plasmid’s molecular weight and Avogadro’s constant (Rizza *et al*., 2009). Regression analysis was adjusted to the standard curves using Excel software (Microsoft). qRT-PCR was performed on four biological and three technical replicates.

### Transmission of recombinant viruses by *Rhopalosiphom maidis*

A sufficient virus free population of *Rhoalosiphum maidis* were maintained at 2-3 hours under starvation condition. It was fed from the inoculated plants with recombinant constructs and wild type viruses for 10-30 minutes. Subsequently, aphids were transferred to the seedling of the healthy Johnsongrass in separate compartments. After appearance of symptoms, leaf samples were verified by RT PCR.

## Acknowledgment

We thank Hannu Pappu for his assistance in reviewing of manuscript.

Authors declare that they have no conflict of interest.

## References

Atreya, C.D., Raccach, B. and Pirone, T.P. 1990. A point mutation in the coat protein abolishes aphid transmissibility of a potyvirus. Virology 178:161–165.

Beelman, D. A., and Parker, R. 1995. Degradation of mRNA in eukaryotes. Cell 81, 179–183.

Gibbs, A. and Ohshima, K., 2010. Potyviruses and the digital revolution. Annual review of phytopathology, 48, pp. 205–223.

Gibson, D.G., 2011. Enzymatic assembly of overlapping DNA fragments. In Methods in enzymology (Vol. 498, pp. 349–361). Academic Press.

Guilford, P. J., Beck, D. L., and Forster, R. L. S. 1991. Influence of the poly(A) tail and putative polyadenylation signal on the infectivity of white clover mosaic potexvirus. Virology 182, 61–67.

Hosseini, A., Habibi, M.K., Izadpanah, K., Mosahebi, G.H., Rubies-Autonell, C. and Ratti, C., 2010. Characterization of a filamentous virus from Bermuda grass and its molecular, serological and biological comparison with Spartina mottle virus. Archives of virology, 155(10), pp. 1675–1680.

Izadpanah, K., M. Zaki-Aghl, Y. P. Zhang, S. D. Daubert and A. Rowhani. 2003. Bermuda grass as a potential reservoir host for *Grapevine fan leaf virus*. Plant Dis. 87:1179–1182

Jiang, J. and Laliberté, J.F., 2011. The genome-linked protein VPg of plant viruses—a protein with many partners. Current opinion in virology, 1(5), pp. 347–354.

Jorda, C., Ortega, A. & Juarez, M. 1995. New hosts of tomato spotted wilt virus. Plant Disease, 79.

Leng, P., Ji, Q., Tao, Y., Ibrahim, R., Pan, G., Xu, M. and Lübberstedt, T., 2015. Characterization of sugarcane mosaic virus Scmv1 and Scmv2 resistance regions by regional association analysis in maize. PloS one, 10(10), p. e0140617.

Lockhart, B., Khaless, N., El Maataoui, M. & Lastra, R. 1985a. Cynodon chlorotic streak virus, a previously undescribed plant rhabdovirus infecting Bermuda grass and maize in the Mediterranean area. Phytopathology, 75, 1094–1098.

Lockhart, B., Khaless, N., Lennon, A. M. & El Maatauoi, M. 1985b. Properties of Bermuda Grass Etched-Line Virus, A New Leafhopper-Transmitted Virus Related to Maize Rayado Fino and Oat Blue Dwarf Viruses. Phytopathology, 75, 1258–1262.

Lopez-Moya, J.J., Wang, R.Y. and Pirone, T.P., 1999. Context of the coat protein DAG motif affects potyvirus transmissibility by aphids. Journal of General Virology, 80(12), pp. 3281–3288.

Louie, R. 1980. Sugarcane msaic virus in Kenya. Plant Dis. 64:994–947.

Mostafavi Neishaburi, F. S., Masumi, M., Nasrollanejad, S., Rahpeyma-Sarvestani, N., & Izadpanah, K. 2015. Analyses of complete nucleotide sequence of Iranian isolate of maize dwarf mosaic virus (MDMV) and notes on the origin and evolution of MDMV. Iranian J. Plant Pathology 51(1):69–81.

Rizza S, Nobile G, Tessitori M Catara A, Conte E, 2009. Real time RT-PCR assay for quantitative detection of *Citrus viroid III* in plant tissues. Plant Pathology 58, 181–185.

Stewart, L.R., Bouchard, R., Redinbaugh, M.G. and Meulia, T., 2012. Complete sequence and development of a full-length infectious clone of an Ohio isolate of Maize dwarf mosaic virus (MDMV). Virus research, 165(2), pp. 219–224.

Tacahashi, Y. and Uyeda, I., 1999. Restoration of the 3′ end of potyvirus RNA derived from poly (A)-deficient infectious cDNA clones. Virology, 265(1), pp. 147–152.

Tatineni, S., McMechan, A.J., Bartels, M., Hein, G.L. and Graybosch, R.A., 2015. In vitro transcripts of wild-type and fluorescent protein-tagged Triticum mosaic virus (family Potyviridae) are biologically active in wheat. Phytopathology, 105(11), pp. 1496–1505.

Uzarowska, A., Dionisio, G., Sarholz, B., Piepho, H. P., Xu, M., Ingvardsen, C. R., Lubberstedt, T. 2009. Validation of candidate genes putatively associated with resistance to SCMV and MDMV in maize (Zea mays L.) by expression profiling. BMC plant biology, 9(1-15).

Wolf, J.B., Künstner, A., Nam, K., Jakobsson, M. and Ellegren, H., 2009. Nonlinear dynamics of nonsynonymous (dN) and synonymous (dS) substitution rates affects inference of selection. Genome biology and evolution, 1, pp. 308–319.

Zare, A., Masumi, M. and Izadpanah, K. 2005. Bermuda grass southern mosaic virus: A distinct potyvirus infecting several gramineous species in Iran. Parasitica 61: 105–110.

